# The shape of gene expression distributions matter: how incorporating distribution shape improves the interpretation of cancer transcriptomic data

**DOI:** 10.1101/572693

**Authors:** Laurence de Torrenté, Samuel Zimmerman, Masako Suzuki, Maximilian Christopeit, John M. Greally, Jessica C. Mar

## Abstract

In genomics, we often impose the assumption that gene expression data follows a specific distribution. However, rarely do we stop to question this assumption or consider its applicability to all genes in the transcriptome. Our study investigated the prevalence of genes with expression distributions that are non-Normal in three different tumor types from the Cancer Genome Atlas (TCGA). Surprisingly, less than 50% of all genes were Normally-distributed, with other distributions including Gamma, Bimodal, Cauchy, and Lognormal were represented. Relevant information about cancer biology was captured by the genes with non-Normal gene expression. When used for classification, the set of non-Normal genes were able to discriminate between cancer patients with poor versus good survival status. Our results highlight the value of studying a gene’s distribution shape to model heterogeneity of transcriptomic data. These insights would have been overlooked when using standard approaches that assume all genes follow the same type of distribution in a patient cohort.

## Introduction

A fundamental tenet of applied statistics states that under certain conditions, data can be modeled by a probability distribution. If appropriately applied, then this assumption is extremely powerful because it represents a fast track to well-established statistical methods to address questions about the data. Like many statistical measures in biology, we often assume that gene expression follows a Normal distribution, and the most common statistical methods, such as the t-test and ANOVA models, are predicated on this assumption. As we begin to learn more about the diversity of gene expression, we call into question the relevance that just one distribution applies to all genes. Our study addresses a simple question – how prevalent is Normality in gene expression and what new information can we learn by going beyond the Normal distribution? We raise this question not to invalidate previous findings that have been based on assumptions of Normality, but instead to draw attention to genes that may have otherwise been overlooked and the insights that they bring, especially in the context of disease processes.

Using the Cancer Genome Atlas (TCGA), we show that greater than 50% of genes in cancer transcriptomes are non-Normally distributed for multiple tumor types. Most significantly, we show that accounting for the distribution shape improved the prediction accuracy of patient survival time. Our study demonstrates that the assumption of Normality does not apply uniformly to all genes in the cancer transcriptome, and in fact, a range of distributions are represented. Importantly, incorporating non-Normality into the analysis of gene expression data revealed information for understanding transcriptional control of cancer that would have been missed using standard approaches.

To investigate the prevalence of Normally-distributed genes in the transcriptome, we assembled a panel of six statistical distributions. Each distribution has their own properties and collectively capture a diversity of density shapes (Figure 1). We included two symmetric distributions, the Normal and Cauchy distributions, where the latter has heavy tails and is more peaked than the Normal distribution. The Lognormal, Pareto and Gamma distributions all have skewed, asymmetric shapes. The Lognormal is an asymmetric distribution but on a log scale, so Normality assumptions are still applicable for this distribution. In contrast, the Pareto is a heavy-tailed distribution and the most skewed on our distribution panel. The Gamma is a distribution whose values can span more or less skewness depending on the parameters, and overall its shape is not as extreme as the Pareto. The Bimodal distribution models an alternative kind of heterogeneity where two distinct sub-groups exist in the data. We applied this panel to three different TCGA datasets, the acute myeloid leukemia (AML) [1], ovarian cancer (OVC) [2] and Glioblastoma multiforme (GBM) [3] patient cohorts.

**Figure 1.**
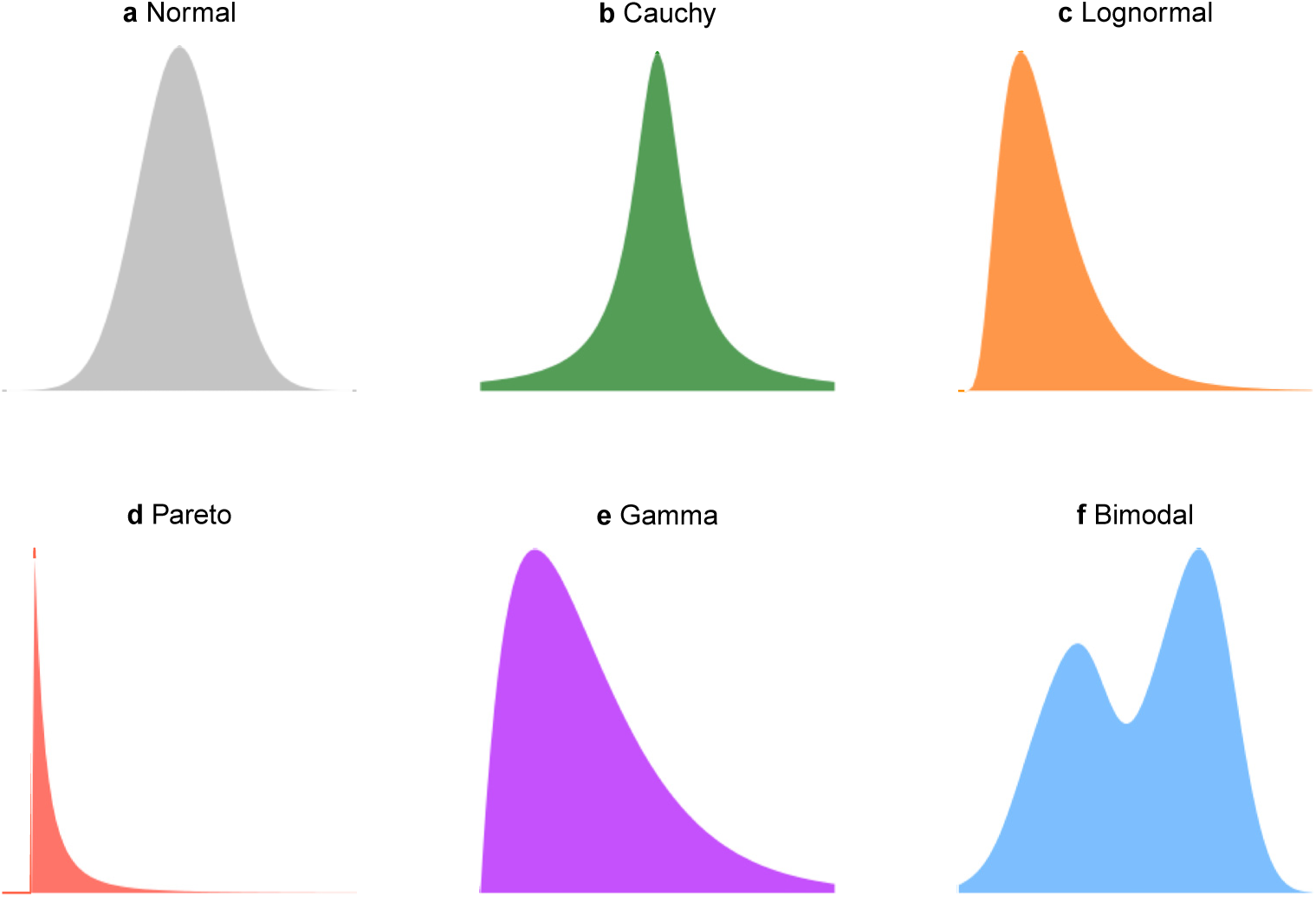
Panel of six statistical distributions that capture a diversity of different probability density shapes. **A.** Normal, **B.** Cauchy, **C.** Lognormal, **D.** Pareto, **E.** Gamma, **F.** Bimodal.

## Results

### Over 50% of genes in the cancer transcriptome does not follow a Normal distribution

A classification scheme was developed to assign genes based on how similar their expression distributions were to one of the six probability density distributions described above (Figure 2). First, the classification scheme was designed to evaluate the statistical likelihood of whether a gene’s expression profile matched a Bimodal distribution, and if this was not the case, to then assess whether any of the remaining five unimodal distributions were a more appropriate fit (see Materials and Methods). If an adequate fit could not be determined from these six options, the gene was discarded from further analysis. Under this classification scheme, the Normal distribution was found to capture only 13 to 15% of genes in the three microarray cancer datasets in this study (13.73% for AML, 13.52% for GBM, 15.13% for OVC, Figure 3A) and less than 45% of genes in the RNA-seq datasets for the same tumor types (30.29% for AML, 41.8% for GBM, 43.18% for OVC, Figure 3C). The Gamma distribution was the largest non-Normal category of genes for both microarray and RNA-seq datasets (representing 21-32%). Since different microarray platforms were used to generate the data, we also investigated whether this could have had any effect on the distribution counts observed (Table S1). When the analysis focused only on the genes that were common across all array platforms, it could be seen that the proportion of genes assigned to each of the distribution remained the same, indicating this effect is more likely biological rather than technical (Figure 3B).

**Figure 2.**
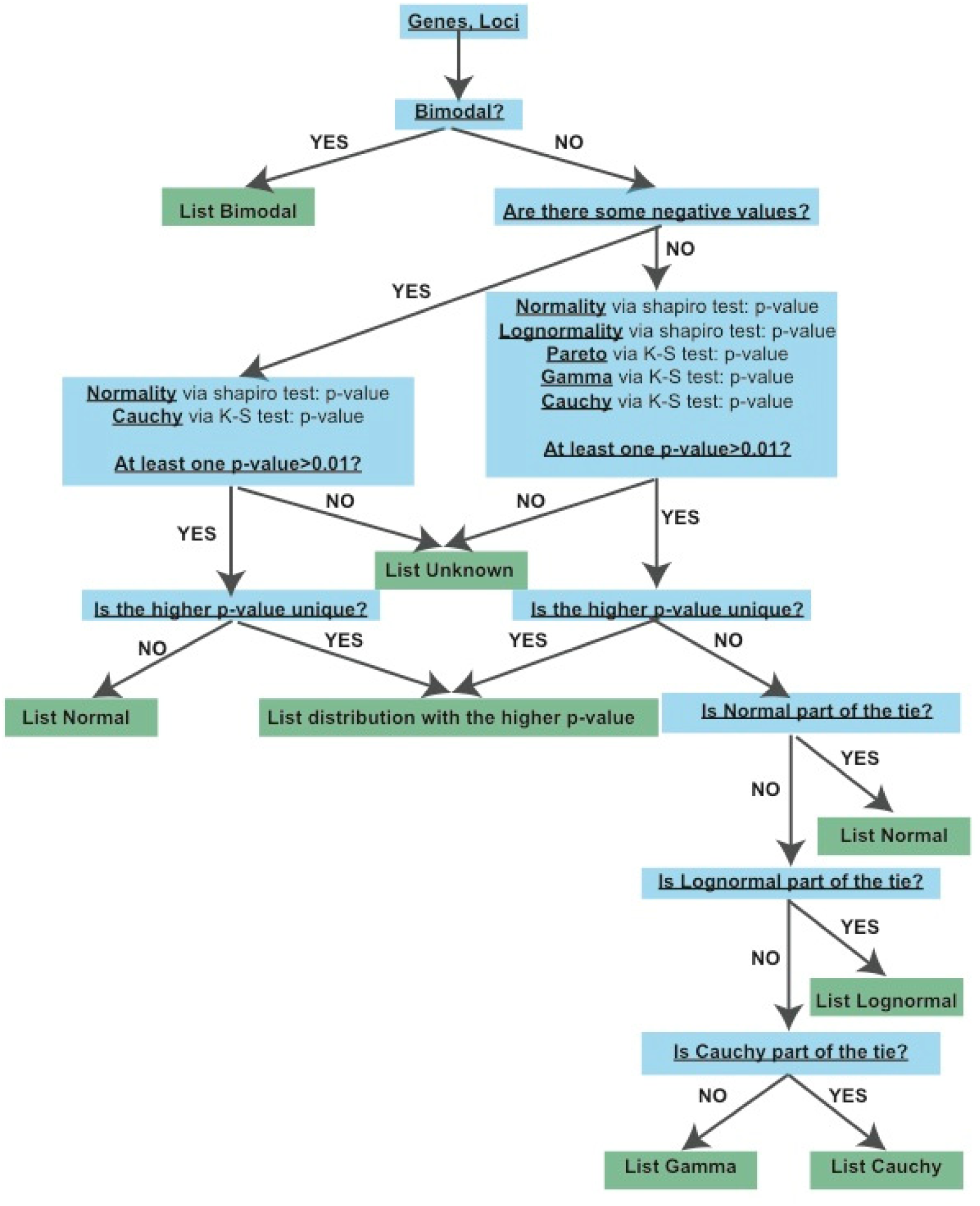
Classifying genes according to their shape of gene expression distribution. If a locus/gene is classified as bimodal, it is removed from the list and the algorithm continues on the remaining loci/genes. The category sets are mutually exclusive. It means that one gene/locus is classified in only one category. Genes/loci that fall in neither category are classified as unknown.

**Figure 3.**
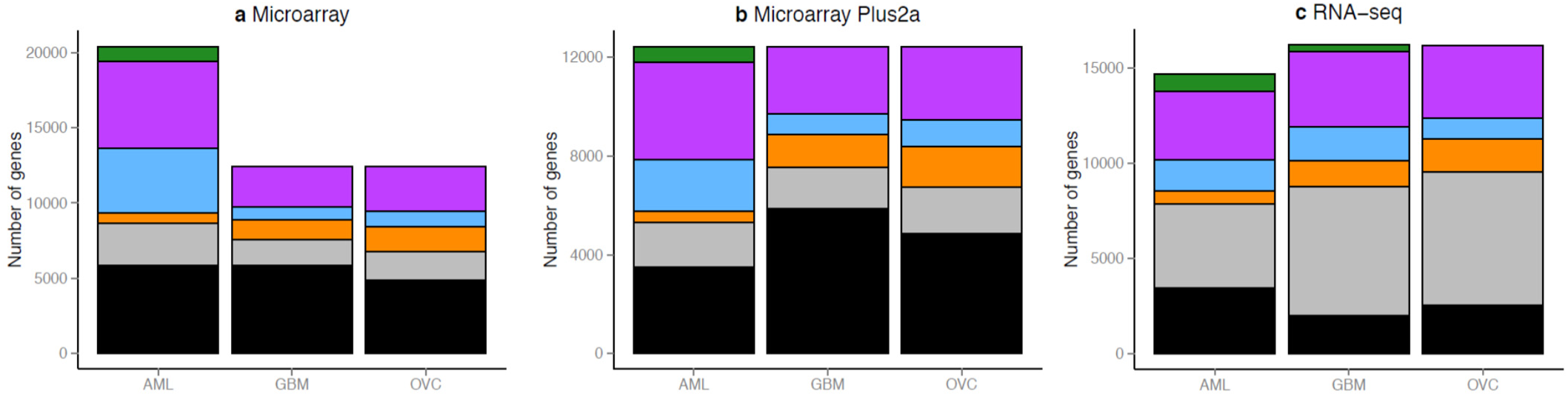
Representation of genes and loci in each distribution category. **A.** Number of genes classified in each distribution for the three microarray datasets, **B.** when exclusively looking at the subset of genes in the Plus2a platform, **C.** for the three RNA-seq datasets.

### Non-Normally expressed genes can discriminate between good versus poor patient survival outcomes

In cancer patient cohorts, gene expression is commonly used to identify genes that can discriminate between patients with good versus poor survival. Typically, the methods employed for this purpose assume that all genes follow the same distribution. This assumption is limited because identifying meaningful sub-groups is dependent on how a gene’s expression profile is distributed in the patient cohort. For example, if a gene is assumed to have Normally-distributed data (or any symmetric distribution), we typically compare the patients that have the expression of a gene in the extreme tails of the distribution, against those with gene expression in the non-tail region (Figure 4A). Because of the symmetry of the data distribution, this is a rational sub-grouping to adopt. However, for other distributions that are non-symmetric, using both tails of the distribution to form one patient group is not an intuitive sub-grouping. Instead, a tail versus non-tail (Figure 4B), or one mode versus the second mode (Figure 4C) is more instructive for asymmetric and bimodal distributions, respectively, for contrasting survival curves (Figure 4D).

**Figure 4.**
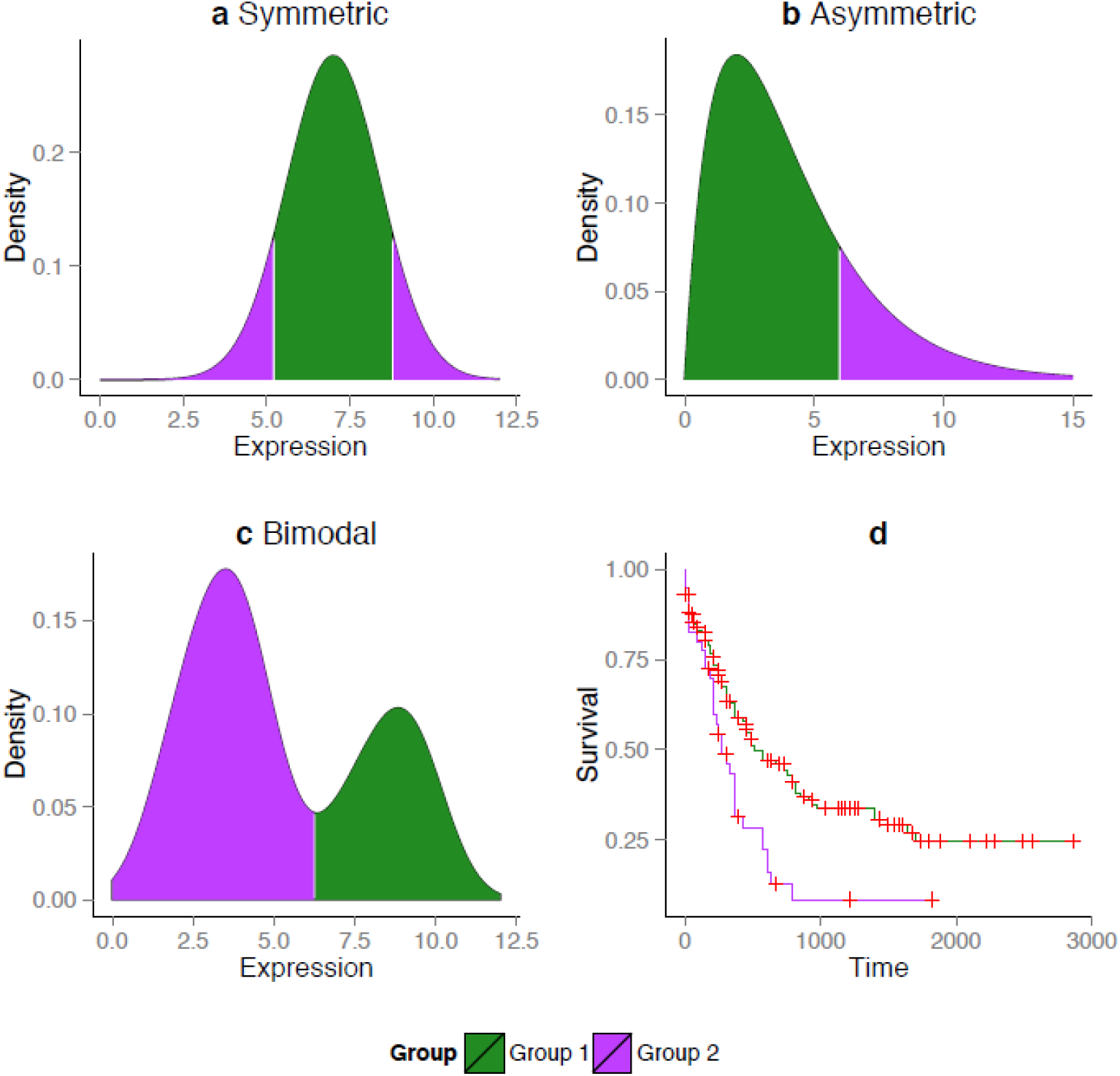
Decision rules for identifying extreme and non-extreme patient groups that take into account the specific shape of the gene expression distribution. **a.** For symmetric distributions, the extreme patient group is represented by the samples falling in either the upper or lower percentile of the distribution as shown by the purple tails (in our analysis, the tenth percentile is used). The non-extreme patient group corresponds to the samples falling between these two percentile cut-offs as shown by the green region. **b.** For asymmetric distributions, the extreme patient group corresponds to samples only in the first or last percentile depending on the shape of the asymmetry, as shown by the one-sided purple tail. The remaining region of the distribution represents the non-extreme patient group. **c.** For bimodal distributions, the split is determined by a clustering algorithm applied to the expression data to identify which patients belong to one group (mode) versus another. For genes in the bimodal expression category, the definition of extreme and non-extreme patient groups is not relevant, and instead we identify two patient groups for comparison, as shown by the purple and green regions. **d.** Theoretical example of two survival curves constructed for patients in Groups 1 and 2 as defined in a, b or c.

In this study, we refer to prognostic marker genes as those genes whose expression distinguishes significant differences in patient survival time. We found that when distribution shape is accounted for, a set of prognostic marker genes was identified that was almost unique compared to when standard assumptions were applied (log-rank test, P-value < 0.05, Table S2). The intersection between genes found when the distribution shape information was used, and those when the assumption of symmetry was applied uniformly, was minimal for the three cancers (for the non-symmetric genes, the overlap was eight genes for AML, and zero for GBM and OVC). This small overlap indicates that a very different set of prognostic markers were identified, depending on whether the distribution shape was factored in.

The second largest type of non-Normal distribution represented amongst the genes that were identified based on the distribution shape information was the Bimodal distribution. This result highlights the existence of Bimodally-expressed genes that have distinct modes corresponding to statistically significant differences in survival time. Genes that were found to be significant for patient survival time using the shape of the expression distribution are listed in Table S3. To determine the degree of robustness of our result, we also looked at how many genes were significant in survival time when patients were randomly assigned to the two groups (corresponding to the green and purple regions in Figure 4). No genes were significant under this assumption, indicating that the genes found using the distribution shape information were unlikely to be detected purely by chance (Table S2).

### Identifying prognostic marker genes using the expression distribution shape information provides functional insights into cancer biology that were not found using standard symmetric assumptions

For AML, functional terms and biological pathways from MSigDB were significantly over-represented in the non-Normal prognostic marker genes, and not in the set of Normally-expressed prognostic marker genes, indicating that these gene sets correspond to different pathways (Table S5-7). For AML, the non-Normal prognostic marker genes were enriched for the KEGG inositol phosphate metabolism pathway. Previous studies have demonstrated a link between this pathway and cancer, where common germline variation in this pathway has been shown to serve as a susceptibility factor [4, 5]. Another KEGG pathway that was enriched was related to Fc gamma R-mediated phagocytosis, a pathway that has previously been shown to be upregulated in HL-60 cells, which is a leukemia cell line [6]. The non-Normal prognostic marker genes in AML were also enriched for an oncogenic signature based on human leukemia cells from a HOXA9 knockdown (Table S5). We investigated whether any of our non-Normal genes that were identified using the distribution shape information had been detected in seven previous studies of AML gene expression [7–12]. Of all seven signatures tested, we observed at most 2 genes out of 561 in the signature, suggesting that the non-Normal genes we have identified may be prognostic markers of AML that have largely been missed by existing analyses (Table S8).

For OVC, the pathways that were exclusively over-represented in the non-Normally expressed genes were enriched for the MicroRNA biogenesis REACTOME pathway. It has been shown that gene sets related to RNase III DROSHA and DICER1 were decreased in ovarian cancer [13]. The non-Normal prognostic marker genes were also enriched in genes defining epithelial-mesenchymal transition which is a critical step for cancer cell invasion and metastasis [14] (Table S7). For GBM, the non-Normal prognostic marker genes were enriched for the immune system and neuronal system pathways from REACTOME, and more broadly for gene sets involved in the regulation of cellular and biological processes (Table S6). Overall, for this tumor type, the pathways over-represented in the list of non-Normal prognostic marker genes were less specialized and with less of a direct link to cancer compared to the other two tumor types.

### Incorporating the shape of the gene expression distribution improved the performance of a classifier’s ability to predict the survival of individual patients in different types of cancers

Prognostic marker genes can be combined to construct a gene expression-based classifier to predict the survival time of new patients. We investigated whether leveraging information about the expression distribution shape resulted in more accurate predictions of patient survival time compared to classifiers that assumed all genes followed a symmetric distribution. To compare the performance of these two sets of predictors, we used a non-parametric classification method, a random survival forest, on genes that were selected based on whether they were significantly different in expression under the two different assumptions.

Because of its ability to distinguish relevant features from irrelevant ones, the random survival forest method [15] is well-suited for high-dimensional problems. It is an efficient computational method that can handle non-linear or complex higher-order interaction effects. Under the algorithm framework, a tree represents a graphical construct that describes the hierarchical relationship between genes. Individual genes are prioritized in the hierarchy based on how well their expression profiles can discriminate patient survival times. A collection of trees, termed a forest, is grown by the algorithm using independent bootstrapping of the original dataset. For each tumor type, two-thirds of the data was designated for classifier training, and the random survival forest method was applied to this training set using 1000 bootstraps. The remaining one-third of the data was used for testing the accuracy of the classifier by predicting the survival status of each patient that the classifier had not yet seen.

Classifiers were constructed under the two assumptions that differed in how the patient subgroups were identified. First, the shape of the gene expression distribution was taken into consideration so that the subgroups were defined as in Figure 4B. Second, all genes were assumed to have symmetric expression distributions (Figure 4A). To test the robustness of the results, the third set of classifiers was also constructed based on a random selection of genes that was equal in size to the number of genes obtained under the shape-based assumption. To ensure that our results were not biased toward a specific combination of patients in the training and test datasets, we randomly divided the data into training and test sets 100 times. The classification procedure was repeated under the three sets of assumptions for the three different tumor types for each of the 100 unique training/test datasets. Performance of the three classifiers was assessed based on misclassification rates observed for the 100 repeats of each tumor type. A prediction was considered misclassified if a patient was predicted to have good survival when in fact the patient’s survival true status was poor, or vice versa (Figure 5).

**Figure 5.**
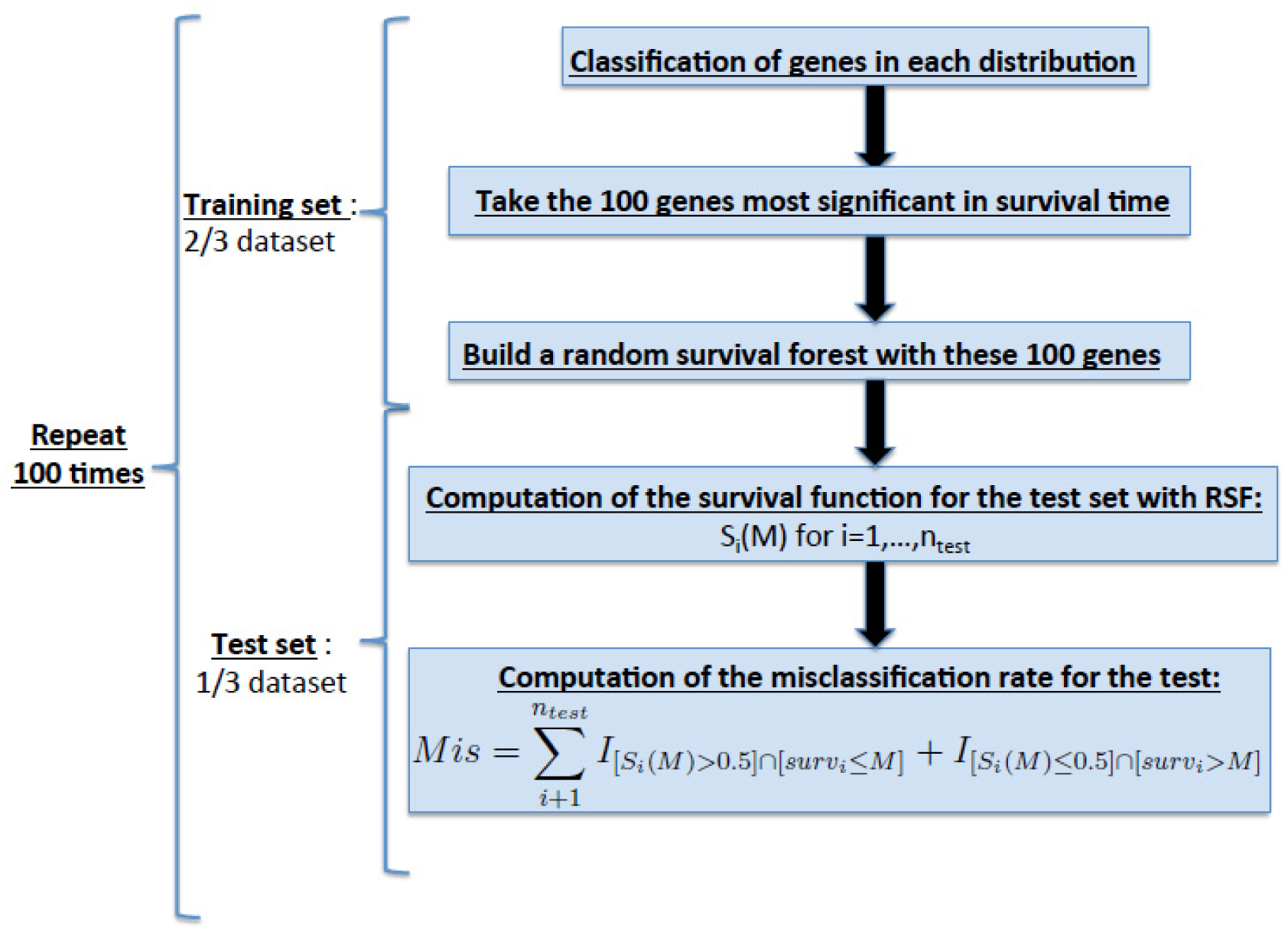
Pipeline to test the performance of the shape-based assumption in patient survival prediction. This pipeline is used once for each assumption: shape, symmetric and random where the difference occurs in step 2. For the shape assumption, we will separate the samples using the shape (Figure 6a-c) and then compute the p-value with a log-rank test. For the symmetric assumption, we will separate the samples into two groups using the same splitting for all the genes, and compute the P-value with a log-rank test. For the random assumption, the 100 genes are just chosen at random from the dataset and then compute the P-value with a log-rank test.

Classifiers derived under the shape-based assumption surpassed the performance of the symmetric-based ones for the microarray and RNA-seq datasets of the three cancer types, AML, GBM and OVC (Figure 6). Performance of the classifiers can also be assessed by counting how many times the shape-based classifier outperformed the symmetric-based classifier in the 100 repeats performed. Using this metric, the shape-based classifier performed as well or better than the symmetric-based one in at least 60/100 repeats (Table 1). In AML, 63/100 for microarray and 60/100, the shape-based classifier had an equal or better misclassification rate than the symmetric one. The two classifiers had an even stronger performance for GBM and OVC (66/100 and 69/100 for microarray, 71/100 and 70/100 for RNA-seq). In summary, building the shape of the gene expression distribution into the prediction of patient survival time results in increased performance.

**Table 1.** Number of times where the shape assumption is better than the symmetric one for Microarray and RNA-seq datasets. The number in parenthesis represent the number of equality: between both misclassification rate. Number of times where the shape assumption is correct compared to the symmetric assumption for patient survival status.

|  | Microarray data |  |  | RNA-seq data |  |  |
| --- | --- | --- | --- | --- | --- | --- |
|  | AML | GBM | OVC | AML | GBM | OVC |
| Misclassification Rate of Shape assumption<br>≤ Misclassification Rate of Symmetric<br>assumption | 63 (7) | 66 (2) | 69<br>(16) | 60 (7) | 71<br>(13) | 70 (14) |

**Figure 6.**
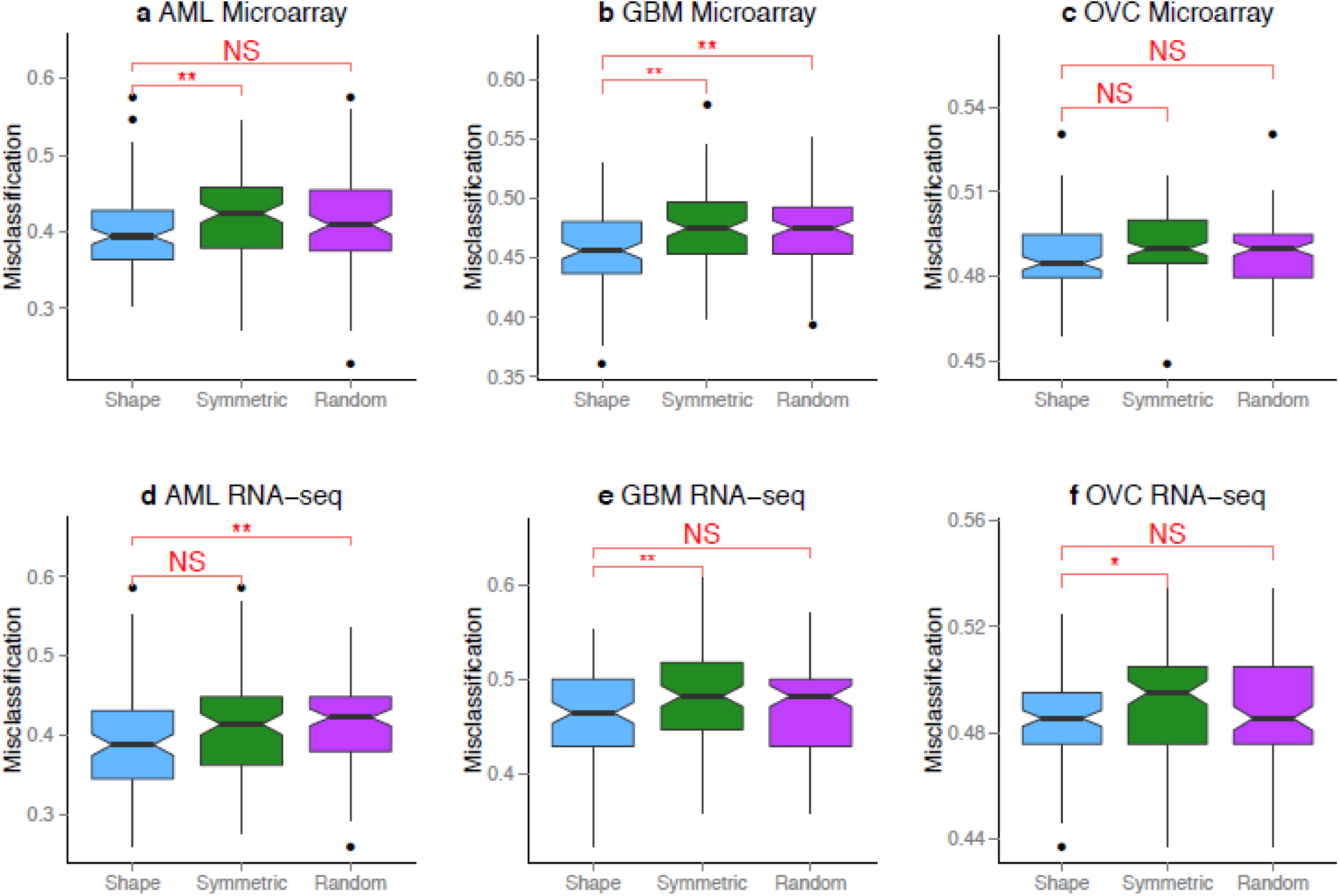
Comparing prediction accuracy using classifiers that incorporate the expression shape versus assuming a symmetric distribution for all genes. We used random survival forests to predict the prognosis of patients and tested the performance of classifiers derived three ways; first, incorporating information from the distribution shape, second, assuming symmetry for all genes, and third,for a random set of genes. Classifiers were trained on 2/3 of the data, tested on 1/3, and repeated 100 times **a** AML Microarray, **b** GBM Microarray, **c** OVC Microarray, **d** AML RNA-seq, **e** GBM RNA-seq, **f** OVC RNA-seq.. Stars indicate datasets where the shape-based approach produced lower misclassification rates that were statistically significant (Wilcoxon test, * = P-value < 0.05, ** = P-value < 0.01, NS = Not Significant). The notch in each boxplot displays a confidence interval based on median misclassification rate±1.58×IQR/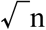 where n = 100, notches that do not overlap reflect statistically significant comparisons.

### Box-Cox transformations did not alter the number of Normally-distributed genes in RNA-seq data

Our results demonstrated an overwhelming representation of non-Normal distributions in the transcriptome, with the range of non-Normally expressed genes being 56.82 to 69.71% in the RNA-seq datasets for all three tumors. In applied statistics, a common procedure to induce Normality for seemingly non-Normal data is a Box-Cox transformation, and one could argue that applying these standard adjustments would restore Normality in the data. To investigate this assumption, the Box-Cox transformations were applied with varying parameters *λ* = −10, ⋯, 10 to both microarray and RNA-seq datasets. For all three RNA-seq datasets, the maximum number of Normally-distributed genes was observed when the Box-Cox transformation was not applied. In other words, application of the Box-Cox transformation was not successful in converting the non-Normally-distributed genes into Normal ones across the parameter space that was used (Figure 6). For the microarray datasets, the number of Normally-distributed genes did increase due to the Box-Cox transformation; however, the difference observed was small (Figure 7).

**Figure 7.**
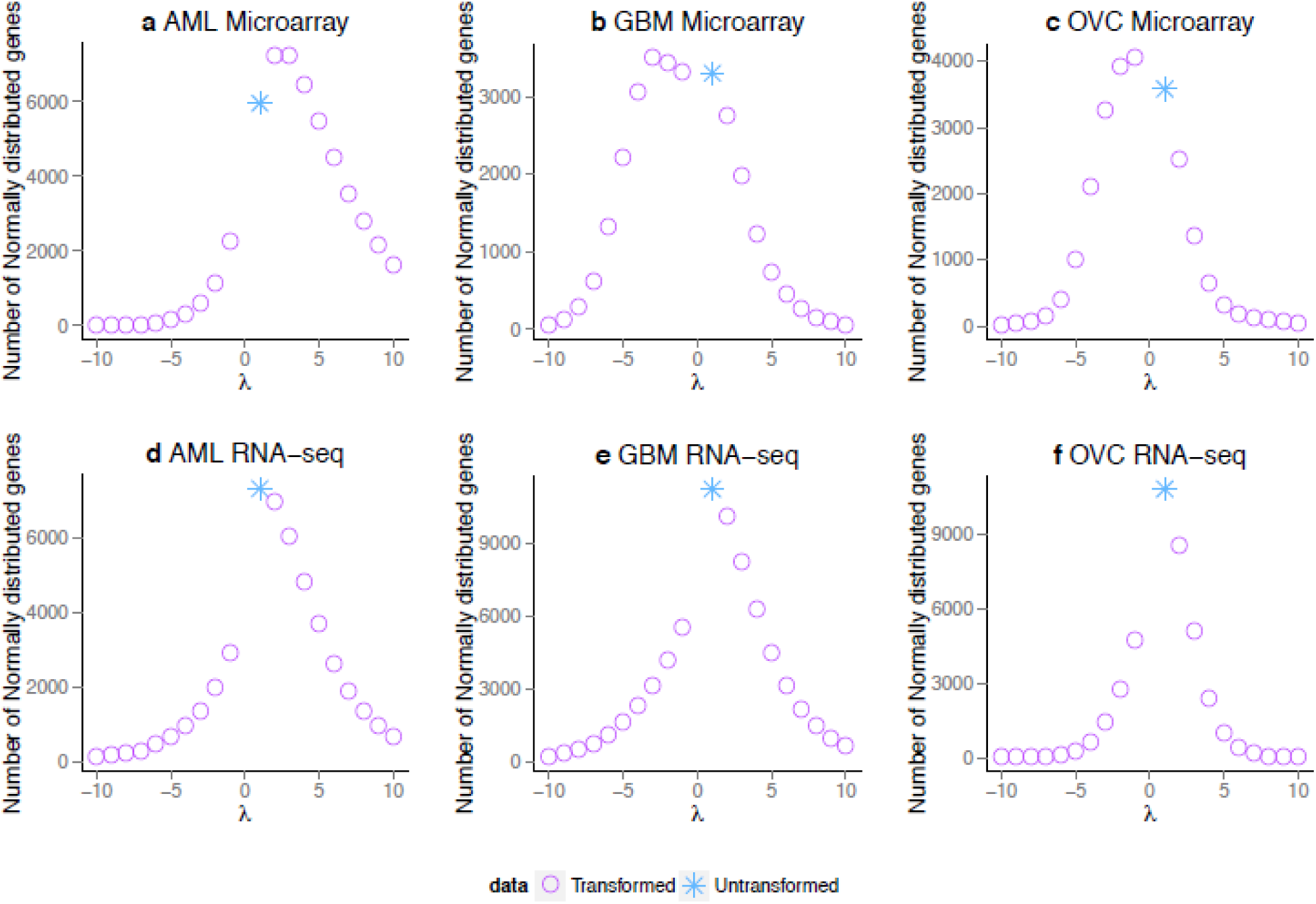
Box-Cox transformation applied to Microarray and RNA-seq datasets. The Box-Cox transformation 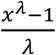 with *λ* = −10, ⋯, 10 was applied to the three microarray and RNa-seq datasets to see if the number of Normally distributed genes was changing. The blue star corresponds to untransformed dataset and the purple circle to the transformed ones.

### Tumor purity does not influence variation in gene expression shape

To assess the influence of tumor purity, we used the information provided by TCGA that represented the pathologist’s estimate of purity for each patient sample and correlated this measure with gene expression. In general, minimal correlation was observed between the microarray, RNA-seq datasets and the tumor purity (Figure S1). For the GBM RNA-seq dataset, no correlations between loci or genes were statistically significant (adjusted P-values < 0.001). For the remaining datasets, the number of significant correlations remained small relative to the total number of genes or loci; where 377, 384 and 441 genes were statistically significant for GBM microarray, OVC microarray and OVC RNA-seq datasets respectively.

## Discussion

Our study illustrates just how diverse distributions can be in cancer transcriptomes. We often take for granted that genes follow the same expression distribution and that it is their population-level summary statistics, like the average gene expression, that will identify key regulators of disease. While these summary statistics are indeed important, we showed that modeling other features of the expression distribution could also provide regulatory information. Fundamentally, the results of our study are significant because they force us to confront the fact that a gene’s expression profile cannot simply be summarized by a single statistical distribution. We showed how incorporation of the shape of the expression distribution identified genes with prognostic value for patient survival status that was not detected using conventional approaches. Moreover, we showed that using this shape information of the expression distribution resulted in a more accurate classification of patient survival time.

Throughout the history of science, the Normal distribution has been a ubiquitous feature in many forms of data analysis. Part of this ubiquity can be attributed to the central limit theorem (CLT), which explains how the average value of a variable will approximately follow a Normal distribution, regardless of the underlying data distribution. The validity of the CLT is dependent upon the data being sufficiently large and having been sampled independently from the same population. Because of the CLT, standard statistical methods typically have some degree of inbuilt robustness so that they are generally able to produce valid inferences even in the presence of some non-Normal data, but this does not apply to all situations. In reality deviations from Normality do exist in the data, but the extent of these deviations is not commonly assessed. For the entire transcriptome, this means that genes with expression profiles that more closely resemble a Normal distribution will be more easily detectable by standard statistical methods. This kind of bias means many genes may be being overlooked or down-weighted because we are not stopping to first evaluate the prevalence of different distributions [16].

Attention to non-Normality in gene expression has so far yielded some valuable insights in cancer biology. For instance, in a patient cohort, genes with distinct on and off transcriptional states followed a Bimodal distribution and have been detected in a variety of different tumor types [17]. These switch-like genes have been shown to identify patients subgroups with different rates of survival [18] or distinguish between extremely aggressive forms of tumors [19]. During the early development of statistical models for cDNA microarrays, Newton et al. [20] adopted a Gamma distribution to estimate the significance of gene expression ratios. Despite the mathematical advantages of using the Gamma distribution, this study observed that the overall fit of the Gamma distribution to the entire transcriptome was relatively poor and the use of the distribution was discarded in favor of other more tractable distributions. If we interpret this finding from a different perspective, it is interesting to note that some genes that Newton et al. [20] surveyed had expression profiles that showed a good fit to a Gamma distribution, while others did not.

The search for other non-Normal distributions in the transcriptome remains limited despite the fact that these distributions have the potential to model rare regulatory events in large patient cohorts with more flexibility than a Normal distribution. Non-Normal distributions that are asymmetric or skewed can more accurately model genes spanning a range of aberrant expression for an extreme group of individuals than a symmetric distribution. Such long-tailed aberrations could reflect DNA sequence or copy number variation, different isoforms or alternative splicing patterns. Non-genetic factors at the environmental or epigenetic level may also drive the appearance of different sub-groupings of gene expression in the patient cohort.

The classification of genes into their respective expression distribution shapes may provide an avenue to integrate data of different genomic types such that regulatory mechanisms can be studied more fruitfully. For example, in the promoter region upstream of a gene with Bimodally-distributed expression, a polymorphism may exist such that patients in one expression mode have this mutation, while patients in the other expression mode do not. Similarly, genes with asymmetric expression distributions may be a product of patients who broadly share the same genomic features with a separate minority of patients whose outlier gene expression values reflect differences in methylation, alternative splicing or other regulatory events. Integration of clinical patient data with other genomic data types based on the shape of the distribution may be a more realistic way to identify significant relationships. This is because summary statistics are derived from the total population, i.e., an average expression assumes that all patients will have approximately, a specified level of gene expression.

Genome sequencing projects like TCGA, but also ICGC, HAPMAP, ENCODE, and 1000 Genomes have given us a deeper appreciation for how heterogeneous human populations are concerning genomic features [21]. In light of this, it seems overly simplistic to assume that all key regulators will be found by correlating different data types on the assumption that all patients in the population will exhibit similar levels of the variable of interest. Instead, a more comprehensive approach may be based on identifying subgroupings of patients that share similar levels of a variable and investigating whether there are correlations with other genomic features. Our study has shown how subgroupings can be identified by considering the shape of the expression distribution, and more generally, this method provides a natural way to model heterogeneity under a statistical framework.

## Materials and Methods

### The Cancer Genome Atlas datasets

Data were sourced from The Cancer Genome Atlas (TCGA, http://cancergenome.nih.gov/). The acute myeloid leukemia (AML) dataset had 197 samples with microarray data, and 173 samples with RNA-seq data. The glioblastoma multiforme (GBM) had 549 samples with expression data, and 169 samples with RNA-seq data. The ovarian serous cystadenocarcinoma (OVC) was used only for expression and RNA-seq data, and had 586 samples and 309 samples respectively. For gene expression, the level 2 data on U133A (with 22277 probes corresponding to 12496 genes) for glioblastoma, ovarian and lung, and U133_plus_2 (with 54613 probes corresponding to 19850 genes) for AML were used in this study. For RNA-seq, for AML, GBM and OVC, we used the level 3 IlluminaHiSeq_RNAseqV2 with a total of 20531 genes. We downloaded the clinical data corresponding to the microarray and RNA-seq datasets. From these files, the survival time and the tumor purity estimated by a pathologist (for GBM and OVC) were used. The missing values in survival time were omitted from all statistical analyses.

### Data preprocessing

Gene expression values were log_2_-transformated. For RNA-seq, the genes were filtered by removing those with more than 25% of the samples with values < 1 on the log_2_-transformed scale. GBM and OVC data were batch-corrected using ComBat from the R package sva (version 3.14.0). With the gene expression dataset, we used the annotation R package hgu133a.db (version 3.1.3) for GBM and OVC and hgu133plus2.db (version 3.1.3) for AML. To resolve multiple probes mapping to a unique gene identifier, probes for the same gene symbol were averaged over their corresponding expression levels.

### Classifying genes and loci into different distribution

For testing Normality and Lognormality, the Shapiro test was used from the R package stats (version 3.2.2) [22] with a threshold of 0.01 on the data and log of the data respectively. For Pareto, Gamma and Cauchy, the Kolmogorov-Smirnov test [23] was used. For this test, we needed to set parameter values. For Pareto and Gamma, the parameters were estimated with the Maximum Likelihood Estimates (MLE). For the MLE of Gamma, we used the rGammaGamma R package (version 1.0.12.). For Cauchy, the two parameters were set as the median and the interquartile range. As we are estimating the parameters directly on the dataset, we applied a parametric bootstrap to estimate the final p-value. This idea of resampling to find the null distribution of the test statistics when estimating the parameters is based on the Lilliefors test [24]. The threshold for the final p-value was set to 0.01 for the significance. For testing Bimodality, we computed the Bimodality Index [25] from the R package ClassDiscovery (version 3.0.0.) and kept every gene and locus having a score bigger than 1.1. To improve the speed of the algorithm, we first test if a locus or gene is bimodal, if yes it is classified as so and removed from the list and otherwise the other distributions are tested and the best was chosen (see Figure 2). As the Lognormal, Pareto and Gamma distribution are defined on positive values, they were tested only on genes or locus having all their values bigger than zero. If we have an equality between two distributions the order of classification is as followed: 1) Normal 2) Lognormal 3) Cauchy. Genes that fall in neither category are classified as unknown and the distributions sets are disjoint.

### Evaluating differences in survival time

In order to test the difference between two survival curves, we used the log-rank test from the R package survival (version 2.38.3) with a threshold of 0.05. To estimate the survival curves Kaplan-Meier estimate was used.

## Author Contributions

LDT and SZ performed the analyses. LDT, MC and JCM designed the analyses with input and direction from MS and JMG. LDT and JCM wrote the manuscript that was approved by all co-authors.

