## Supplemental Info for "The shape of gene expression distributions matter: how incorporating distribution shape improves the interpretation of cancer transcriptomic data"

**Supplementary Info**

**Figure S1. Correlation between tumor purity and the microarray, RNA-seq and methylation datasets.** We computed the Spearman correlation between the pathologist estimates of tumor purity and the expression or beta values.These histograms represent the correlation coefficients with, in pink, the coefficients that were significant (p-values<0.01). **a** GBM Microarray with 377 significant coefficients, **b** OVC Microarray with 384 significant coefficients, **c** GBM RNA-seq with no significant coefficients, **d** OVC RNA-seq with 441 significant coefficients.

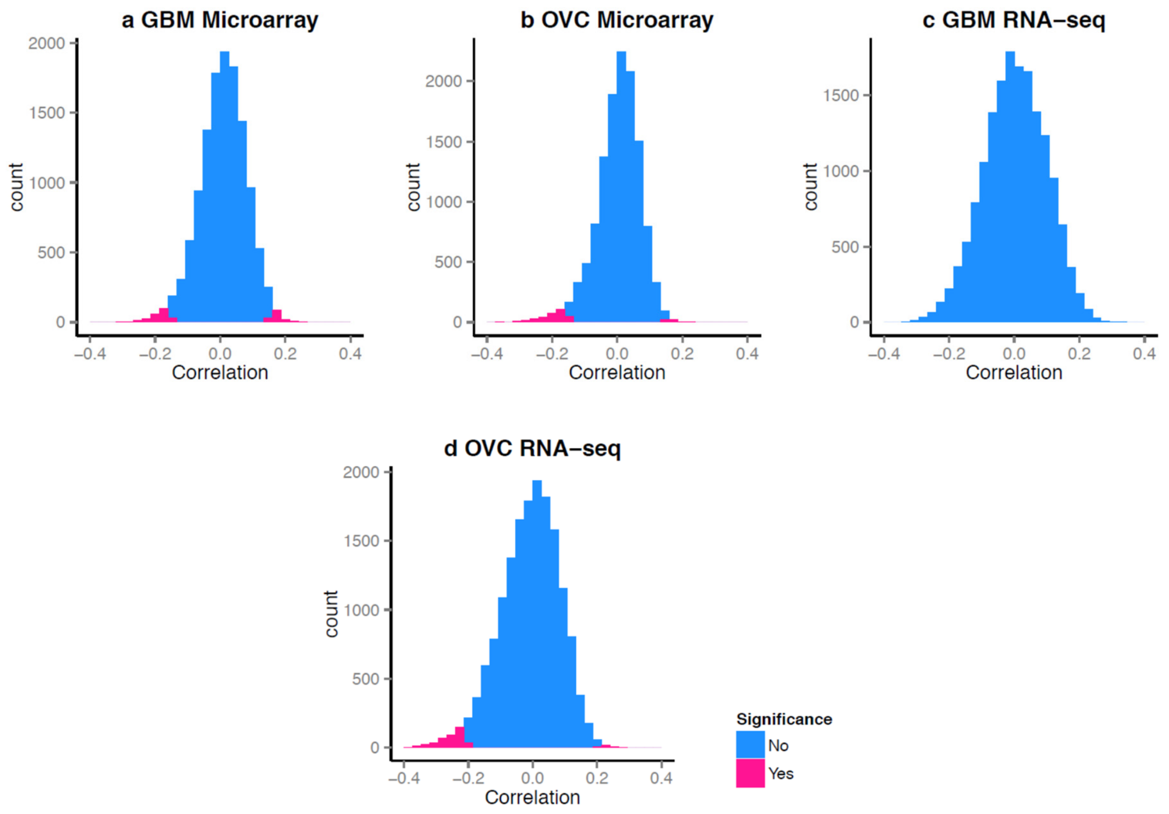

**Table S1. Number of genes and loci in each distribution for AML, GBM and OVC.** For the microarray dataset, AML was done on a hgu133plus2 platform which contains more probes than the hgu133plus2A platform that was used for GBM and OVC. To have a better comparison, we decided to report also the number corresponding to the intersection with the plus2A platform for AML (cf. AML (plus2A)).

| | Bimodal | Normal | Lognormal | Gamma | Cauchy | Pareto | Unknown | Total |
|---|---|---|---|---|---|---|---|---|
| **Microarray** | | | | | | | | |
| AML | 4,323 (21.20%) | 2,799 (13.73%) | 679 (3.33%) | 5,779 (28.34%) | 958 (4.70%) | 0 (0%) | 5,851 (28.70%) | 20,389 |
| AML (plus2A) | 2,113 (16.99%) | 1,809 (14.55%) | 451 (3.63%) | 3,925 (31.57%) | 637 (5.12%) | 0 (0%) | 3,499 (28.14%) | 12,434 |
| GBM | 847 (6.81%) | 1,681 (13.52%) | 1,319 (10.61%) | 2,714 (21.83%) | 1 (0.01%) | 0 (0%) | 5,872 (47.23%) | 12,434 |
| OVC | 1,065 (8.57%) | 1,881 (15.13%) | 1,653 (13.29%) | 2,973 (23.91%) | 0 (0%) | 0 (0%) | 4,862 (39.10%) | 12,434 |
| **RNA-seq** | | | | | | | | |
| AML | 1,637 (11.15%) | 4,447 (30.29%) | 683 (4.65%) | 3,591 (24.46%) | 899 (6.12%) | 0 (0%) | 3,424 (23.32%) | 14,681 |
| GBM | 1,747 (10.77%) | 6,779 (41.80%) | 1,391 (8.58%) | 3,975 (24.51%) | 341 (2.10%) | 0 (0%) | 1,983 (12.23%) | 16,216 |
| OVC | 1,081 (6.68%) | 6,990 (43.18%) | 1,717 (10.61%) | 3,841 (23.73%) | 4 (0.02%) | 0 (0%) | 2,554 (15.78%) | 16,187 |

**Table S2. Number of genes significant in survival time with expression data for a. Microarray and b. RNA-seq datasets.** The total number corresponds to the number of genes in each distribution for each expression data sets. The significance in survival time was assessed using the Log-rank test and a P-value < 0.05. The shape assumption classifies the five distributions into three classes: symmetric (Normal and Cauchy), Bimodal and asymmetric (Gamma and Lognormal). The symmetric assumption uses the symmetric method regardless of the specific distribution. The random assumption takes 20% of the samples in one group and compares them to the remaining 80%. In the second and third columns, the number in brackets corresponds to the number of genes that were found to be significant under both the shape and symmetric assumptions, and the shape and random assumptions, respectively.

**a** Microarray

| | Significant in Survival Time | | | Total Number of Genes |
|---|---|---|---|---|
| | With shape assumption | With symmetric assumption | With a random assumption | |
| **Normal** | | | | |
| AML | 2 | 0 | 0 | 2,799 |
| GBM | 6 | 1 | 0 | 1,681 |
| OVC | 0 | 0 | 0 | 1,881 |
| **Cauchy** | | | | |

| | | | | |
|---|---|---|---|---|
| AML | 0 | 0 | 0 | 958 |
| GBM | 0 | 0 | 0 | 1 |
| OVC | 0 | 0 | 0 | 0 |
| **Lognormal** | | | | |
| AML | 0 | 0 | 0 | 679 |
| GBM | 14 | 0 | 0 | 1,319 |
| OVC | 2 | 0 | 0 | 1,653 |
| **Bimodal** | | | | |
| AML | 7 | 0 | 0 | 4,323 |
| GBM | 30 | 1 | 0 | 847 |
| OVC | 0 | 0 | 0 | 1,065 |
| **Gamma** | | | | |
| AML | 12 | 3 | 0 | 5,779 |
| GBM | 27 | 2 | 0 | 2,714 |
| OVC | 0 | 0 | 0 | 2,973 |
| **Number of Significant Genes** | | | | |
| AML | 21 | 3 | 0 | |
| GBM | 77 | 4 | 0 | |
| OVC | 2 | 0 | 0 | |

**b** RNA-seq

| | **Significant in Survival Time** | | | **Total Number of Genes** |
|---|---|---|---|---|
| | **With shape assumption** | **With symmetric assumption** | **With a random assumption** | |
| **Normal** | | | | |
| AML | 9 | 5 | 0 | 4,447 |
| GBM | 0 | 0 | 0 | 6,779 |
| OVC | 3 | 3 | 0 | 6,990 |
| **Cauchy** | | | | |
| AML | 0 | 0 | 0 | 899 |
| GBM | 0 | 0 | 0 | 341 |
| OVC | 0 | 0 | 0 | 4 |
| **Lognormal** | | | | |
| AML | 4 | 0 | 0 | 683 |
| GBM | 0 | 0 | 0 | 1,391 |
| OVC | 0 | 1 | 0 | 1,717 |
| **Bimodal** | | | | |
| AML | 8 | 1 | 0 | 1,637 |
| GBM | 1 | 0 | 0 | 1,747 |
| OVC | 0 | 0 | 0 | 1,081 |
| **Gamma** | | | | |
| AML | 20 | 0 | 0 | 3,591 |
| GBM | 1 | 20 | 0 | 2,714 |
| OVC | 0 | 0 | 0 | 3,841 |
| **Number of Significant Genes** | | | | |
| AML | 41 | 6 | 0 | |
| GBM | 2 | 0 | 0 | |
| OVC | 3 | 4 | 0 | |

**Table S3. List of genes significant in survival time using the shape assumption per distribution in a. AML Microarray, b. GBM Microarray, c. OVC Microarray, d. AML RNA-seq, e. GBM RNA-seq and f. OVC RNA-seq.**

**a. AML Microarray**

| Distribution | Number of genes | List of genes |
|---|---|---|
| **Normal** | 2 | GPRC6A, IGFBP7 |
| **Bimodal** | 7 | ADAM22, DEGS1, FAM155B, IMPACT, MOB2<br>MSMB, VWC2 |
| **Gamma** | 12 | ABCB11, ASAP1, CRYGA, CSHL1, DCN, EPOR,<br>GPCPD1, INPP5A, LAT, SLC38A5, SPDYE2,<br>TRIM35 |

**b. GBM Microarray**

| Distribution | Number of genes | List of genes |
|---|---|---|
| **Normal** | 6 | KLF9, MMP15, MYLPF, PSMF1, SLC31A2, STX12 |
| **Lognormal** | 14 | APEX2, BRD7, COX4I1, CTCF, EIF3F, ETFB, GATA2<br>HAUS3, IRF2BP1, NOTCH2, PGLYRP4<br>RALYL, TEAD3, XPNPEP3, |
| **Bimodal** | 30 | ALOX12P2, APBA3, BDKRB1, CACNG2, CBFA2T2<br>CLDN5, CLK4, CRCP, DFNA5, EPB41L1, GDF2 |

| | | |
|---|---|---|
| | | GNRHR, GTF3A, IFNAR2, KCNMB1, KLHDC4 LOC79160, MAP1LC3C, METTL1, PPCS, RALBP1 RARG, RBFOX1, RND1, SLC6A15, TAB1, TREML2 TRIM36, UBAP2L, XRCC3 |
| **Gamma** | 27 | AHSG, ALDOB, ARHGEF40, CAPN15, CENPJ, EIF3M GADD45A, LRP5, MAP3K12, MED22, MEF2C, MPZL1 NCR3, PECR, PPP2R5A, PRLR, RANBP10, RPL24 SAP30, SEC11A, SEPP1, SEPT07, SMNDC1, SYK TRIM28, TRPC6, TSPYL5 |

**c. OVC Microarray**

| Distribution | Number of genes | List of genes |
|---|---|---|
| **Lognormal** | 2 | HNRNPF, ZFHX2 |

**d. AML RNA-seq**

| Distribution | Number of genes | List of genes |
|---|---|---|
| **Normal** | 9 | CDCA2, ECE1, KIAA1462, LTB, RGL1, RNASE1 SKA1, SLC1A3, SYNPO2L |
| **Lognormal** | 4 | DUSP7, MTCH1, PPAPDC1B, SLC1A5 |
| **Bimodal** | 8 | CD52, CPNE8, DOCK1, KIAA1958, LOC100130264 RHOBTB3, SPATS2L, SYTL4 |

| | | |
|---|---|---|
| **Gamma** | 20 | ADSS, CAP1, CLINT1, CUX1, DKFZP586I1420<br>GALNT7, GUSBP3, HMMR, ITFG1, LARP1<br>NAP1L1, PNPLA6, RUSC1, SCAMP1, SMAD2<br>SUDS3, TLK2, TUBGCP3, UBE2C, UBQLN1 |

**e. GBM RNA-seq**

| Distribution | Number of genes | List of genes |
|---|---|---|
| **Bimodal** | 1 | RPL39L |
| **Gamma** | 1 | PODNL1 |

**f. OVC RNA-seq**

| Distribution | Number of genes | List of genes |
|---|---|---|
| **Normal** | 3 | C11orf51, CXCL12, PPP1R13L |

**Table S4. Summary of the survival time for a. Microarray and b. RNA-seq.** The survival time is defined as the days to death for the samples that are dead and days to last follow up for the ones still alive. The number of NA relates to the number of samples when the days to death and the last follow up is unknown.

**a Microarray**

| | AML | GBM | OV |
|---|---|---|---|
| **Vital status** | | | |
| Dead | 131 | 462 | 306 |
| Alive | 66 | 86 | 280 |

| | | | |
|---|---|---|---|
| **Median time in days** | 365 | 361 | 896 |
| **Number of NA** | 11 | 4 | 6 |

**b RNA-seq**

| | **AML** | **GBM** | **OV** |
|---|---|---|---|
| **Vital status** | | | |
| Dead | 114 | 131 | 173 |
| Alive | 59 | 37 | 136 |
| **Median time in days** | 335 | 334 | 883 |
| **Number of NA** | 10 | 2 | 2 |

**Table S5. Functional enrichment analyses for genes whose expression distributions are associated with significant patient survival time for the TCGA AML cohort.** Using the Investigate Gene Sets tool from GSEA/MSigDB, we detected significantly enriched terms and pathways for the top 100 most significant in survival time genes using the shape assumption (**a.** Microarray and **c.** RNA-seq), and top 100 most significant in survival time genes using a regular symmetric assumption (**b.** Microarray and **d.** RNA-seq). We used a threshold of FDR q-value < 0.05 or top 10 (whichever is less) was applied, and we computed overlaps with the following sets, H: Hallmark gene sets, C2 : KEGG and REACTOME, C5 : GO biological process, C6 : oncogenic signatures. Blue cells denote terms that were observed for both shape and symmetric assumptions, yellow cells denote terms that were unique to the shape or symmetric assumption (comparison between **a.** and **b.** and between **c.** and **d.**).

**a** Microarray Shape assumption

| Gene Set Name | Description | # Genes in Overlap (k) | # Genes in Gene Set (K) | k/K | p-value | FDR q-value |
|---|---|---|---|---|---|---|
| SYSTEM_DEVELOPMENT | Genes annotated by the GO term GO:0048731. The process whose specific outcome is the progression of an organismal system over time, from its formation to the mature structure. A system is a regularly interacting or interdependent group of organs or tissues that work together to carry out a given biological process. | 10 | 861 | 0.0116 | 4.00E-06 | 7.70E-03 |
| ANATOMICAL_STRUCTURE_DEVELOPMENT | Genes annotated by the GO term GO:0048856. The biological process whose specific outcome is the progression of an anatomical structure from an initial condition to its mature state. This process begins with the formation of the structure and ends with the mature structure, whatever form that may be including its natural destruction. An anatomical structure is any biological entity that occupies space and is distinguished from its surroundings. Anatomical structures can be macroscopic such as a carpel, or microscopic such as an acrosome. | 10 | 1013 | 0.0099 | 1.64E-05 | 1.22E-02 |
| MULTICELLULAR_ORGANISMAL_DEVELOPMENT | Genes annotated by the GO term GO:0007275. The biological process whose specific outcome is the progression of an organism over time from an initial condition (e.g. a zygote or a young adult) to a later condition (e.g. a multicellular animal or an aged adult). | 10 | 1049 | 0.0095 | 2.21E-05 | 1.22E-02 |
| REACTOME_GPVI_MEDIATED_ACTIVATION_CASCADE | Genes involved in GPVI-mediated activation cascade | 3 | 31 | 0.0968 | 2.55E-05 | 1.22E-02 |
| KEGG_FC_GAMMA_R_MEDIATED_PHAGOCYTOSIS | Fc gamma R-mediated phagocytosis | 4 | 97 | 0.0412 | 3.16E-05 | 1.22E-02 |
| REACTOME_SIGNALING_BY_RHO_GTPASES | Genes involved in Signaling by Rho GTPases | 4 | 113 | 0.0354 | 5.75E-05 | 1.84E-02 |
| NERVOUS_SYSTEM_DEVELOPMENT | Genes annotated by the GO term GO:0007399. The process whose specific outcome is the progression of nervous tissue over time, from its formation to its mature state. | 6 | 385 | 0.0156 | 7.79E-05 | 1.87E-02 |
| CENTRAL_NERVOUS_SYSTEM_DEVELOPMENT | Genes annotated by the GO term GO:0007417. The process whose specific outcome is the progression of the central nervous system over time, from its formation to the mature structure. The central nervous system is the core nervous system that serves an integrating and coordinating function. In vertebrates it consists of the brain, spinal cord and spinal nerves. In those invertebrates with a central nervous system it typically consists of a brain, cerebral ganglia and a nerve cord. | 4 | 123 | 0.0325 | 7.99E-05 | 1.87E-02 |
| ORGAN_DEVELOPMENT | Genes annotated by the GO term GO:0048513. Development of a tissue or tissues that work together to perform a specific function or functions. Development pertains to the process whose specific outcome is the progression of a structure over time, from its formation to the mature structure. Organs are commonly observed as visibly distinct structures, but may also exist as loosely associated clusters of cells that work together to perform a specific function or functions. | 7 | 571 | 0.0123 | 8.74E-05 | 1.87E-02 |
| KEGG_INOSITOL_PHOSPHATE_METABOLISM | Inositol phosphate metabolism | 3 | 54 | 0.0556 | 1.37E-04 | 2.63E-02 |

### **b** Microarray Symmetric assumption

| Gene Set Name | Description | # Genes in Overlap (k) | # Genes in Gene Set (K) | k/K | p-value | FDR q-value |
|---|---|---|---|---|---|---|
| SYSTEM_PROCESS | Genes annotated by the GO term GO:0003008. A biological process, occurring at the level of an organ system pertinent to the function of the organism. An organ system is a regularly interacting or interdependent group of organs or tissues that work together to carry out a given biological process. | 8 | 563 | 0.0142 | 1.55E-05 | 1.71E-02 |
| SIGNAL_TRANSDUCTION | Genes annotated by the GO term GO:0007165. The cascade of processes by which a signal interacts with a receptor, causing a change in the level or activity of a second messenger or other downstream target, and ultimately effecting a change in the functioning of the cell. | 13 | 1634 | 0.008 | 1.78E-05 | 1.71E-02 |

**Table S6. Functional enrichment analyses for genes whose expression distributions are associated with significant patient survival time for the TCGA GBM cohort.** Using the Investigate Gene Sets tool from GSEA/MSigDB, we detected significantly enriched terms and pathways for the top 100 most significant in survival time genes using the shape assumption (**a.** Microarray and **c.** RNA-seq), and top 100 most significant in survival time genes using a regular symmetric assumption (**b.** Microarray and **d.** RNA-seq). We used a threshold of FDR q-value < 0.05 or top 10 (whichever is less) was applied, and we computed overlaps with the following sets, H: Hallmark gene sets, C2 : KEGG and REACTOME, C5 : GO biological process, C6 : oncogenic signatures. Blue cells denote terms that were observed for both shape and symmetric assumptions, yellow cells denote terms that were unique to the shape or symmetric assumption (comparison between **a.** and **b.** and between **c.** and **d.**).

**a** Microarray Shape assumption

| Gene Set Name | Description | # Genes in Overlap (k) | # Genes in Gene Set (K) | k/K | p-value | FDR q-value |
|---|---|---|---|---|---|---|
| MULTICELLULAR_ORGANISMAL_DEVELOPMENT | Genes annotated by the GO term GO:0007275. The biological process whose specific outcome is the progression of an organism over time from an initial condition (e.g. a zygote or a young adult) to a later condition (e.g. a multicellular animal or an aged adult). | 12 | 1049 | 0.0114 | 2.02E-06 | 2.10E-03 |
| BIOPOLYMER_METABOLIC_PROCESS | Genes annotated by the GO term GO:0043283. The chemical reactions and pathways involving biopolymers, long, repeating chains of monomers found in nature e.g. polysaccharides and proteins. | 15 | 1684 | 0.0089 | 2.18E-06 | 2.10E-03 |
| REACTOME_NEURONAL_SYSTEM | Genes involved in Neuronal System | 6 | 279 | 0.0215 | 2.78E-05 | 1.79E-02 |
| PROTEIN_METABOLIC_PROCESS | Genes annotated by the GO term GO:0019538. The chemical reactions and pathways involving a specific protein, rather than of proteins in general. Includes protein modification. | 11 | 1231 | 0.0089 | 5.38E-05 | 2.22E-02 |
| NUCLEOBASENUCLEOSIDENUCLEOTIDE_AND_NUCLEIC_ACID_METABOLIC_PROCESS | Genes annotated by the GO term GO:0006139. The chemical reactions and pathways involving nucleobases, nucleosides, nucleotides and nucleic acids. | 11 | 1244 | 0.0088 | 5.91E-05 | 2.22E-02 |
| RNA_METABOLIC_PROCESS | Genes annotated by the GO term GO:0016070. The chemical reactions and pathways involving RNA, ribonucleic acid, one of the two main type of nucleic acid, consisting of a long, unbranched macromolecule formed from ribonucleotides joined in 3',5'-phosphodiester linkage. | 9 | 841 | 0.0107 | 6.92E-05 | 2.22E-02 |
| POSITIVE_REGULATION_OF_CELLULAR_PROCESS | Genes annotated by the GO term GO:0048522. Any process that activates or increases the frequency, rate or extent of cellular processes, those that are carried out at the cellular level, but are not necessarily restricted to a single cell. For example, cell communication occurs among more than one cell, but occurs at the cellular level. | 8 | 668 | 0.012 | 8.24E-05 | 2.27E-02 |
| POSITIVE_REGULATION_OF_BIOLOGICAL_PROCESS | Genes annotated by the GO term GO:0048518. Any process that activates or increases the frequency, rate or extent of a biological process. Biological processes are regulated by many means; examples include the control of gene expression, protein modification or interaction with a protein or substrate molecule. | 8 | 709 | 0.0113 | 1.24E-04 | 2.99E-02 |
| REACTOME_IMMUNE_SYSTEM | Genes involved in Immune System | 9 | 933 | 0.0096 | 1.51E-04 | 3.23E-02 |
| REACTOME_DEVELOPMENTAL_BIOLOGY | Genes involved in Developmental Biology | 6 | 396 | 0.0152 | 1.90E-04 | 3.65E-02 |

**b** Microarray Symmetric assumption

| Gene Set Name | Description | # Genes in Overlap (k) | # Genes in Gene Set (K) | k/K | p-value | FDR q-value |
|---|---|---|---|---|---|---|
| ESTABLISHMENT_OF_LOCALIZATION | Genes annotated by the GO term GO:0051234. The directed movement of a cell, substance or cellular entity, such as a protein complex or organelle, to a specific location. | 11 | 870 | 0.0126 | 2.15E-06 | 4.06E-03 |
| HALLMARK_TNFA_SIGNALING_VIA_NFKB | Genes regulated by NF-kB in response to TNF [GeneID=7124]. | 6 | 200 | 0.03 | 4.22E-06 | 4.06E-03 |
| CELL_PROLIFERATION_GO_0008283 | Genes annotated by the GO term GO:0008283. The multiplication or reproduction of cells, resulting in the expansion of a cell population. | 8 | 513 | 0.0156 | 1.28E-05 | 8.22E-03 |
| STK33_UP | Genes up-regulated in NOMO-1 and SKM-1 cells (AML) after knockdown of STK33 [Gene ID=65975] by RNAi. | 6 | 293 | 0.0205 | 3.66E-05 | 1.57E-02 |
| TRANSPORT | Genes annotated by the GO term GO:0006810. The directed movement of substances (such as macromolecules, small molecules, ions) into, out of, within or between cells. | 9 | 795 | 0.0113 | 4.50E-05 | 1.57E-02 |
| ANATOMICAL_STRUCTURE_DEVELOPMENT | Genes annotated by the GO term GO:0048856. The biological process whose specific outcome is the progression of an anatomical structure from an initial condition to its mature state. This process begins with the formation of the structure and ends with the mature structure, whatever form that may be including its natural destruction. An anatomical structure is any biological entity that occupies space and is distinguished from its surroundings. Anatomical structures can be macroscopic such as a carpel, or microscopic such as an acrosome. | 10 | 1013 | 0.0099 | 5.28E-05 | 1.57E-02 |
| PRC2_EDD_UP.V1_DN | Genes down-regulated in TIG3 cells (fibroblasts) upon knockdown of EED [Gene ID=8726] gene. | 5 | 194 | 0.0258 | 5.70E-05 | 1.57E-02 |
| BCAT.100_UP.V1_UP | Genes up-regulated in HEK293 cells (kidney fibroblasts) expressing constitutively active form of CTNNB1 [Gene ID=1499] gene. | 3 | 49 | 0.0612 | 1.52E-04 | 3.65E-02 |
| SYSTEM_PROCESS | Genes annotated by the GO term GO:0003008. A biological process, occurring at the level of an organ system pertinent to the function of the organism. An organ system is a regularly interacting or interdependent group of organs or tissues that work together to carry out a given biological process. | 7 | 563 | 0.0124 | 1.86E-04 | 3.65E-02 |
| REACTOME_DEVELOPMENTAL_BIOLOGY | Genes involved in Developmental Biology | 6 | 396 | 0.0152 | 1.90E-04 | 3.65E-02 |

**c** RNA-seq Shape assumption

| Gene Set Name | Description | # Genes in Overlap (k) | # Genes in Gene Set (K) | k/K | p-value | FDR q-value |
|---|---|---|---|---|---|---|
| POSITIVE_REGULATION_OF_CELLULAR_PROCESS | Genes annotated by the GO term GO:0048522. Any process that activates or increases the frequency, rate or extent of cellular processes, those that are carried out at the cellular level, but are not necessarily restricted to a single cell. For example, cell communication occurs among more than one cell, but occurs at the cellular level. | 9 | 668 | 0.0135 | 1.49E-05 | 1.64E-02 |
| POSITIVE_REGULATION_OF_BIOLOGICAL_PROCESS | Genes annotated by the GO term GO:0048518. Any process that activates or increases the frequency, rate or extent of a biological process. Biological processes are regulated by many means; examples include the control of gene expression, protein modification or interaction with a protein or substrate molecule. | 9 | 709 | 0.0127 | 2.37E-05 | 1.64E-02 |
| ORGAN_DEVELOPMENT | Genes annotated by the GO term GO:0048513. Development of a tissue or tissues that work together to perform a specific function or functions. Development pertains to the process whose specific outcome is the progression of a structure over time, from its formation to the mature structure. Organs are commonly observed as visibly distinct structures, but may also exist as loosely associated clusters of cells that work together to perform a specific function or functions. | 8 | 571 | 0.014 | 3.44E-05 | 1.64E-02 |
| CELL_DEVELOPMENT | Genes annotated by the GO term GO:0048468. The process whose specific outcome is the progression of the cell over time, from its formation to the mature structure. Cell development does not include the steps involved in committing a cell to a specific fate. | 8 | 577 | 0.0139 | 3.70E-05 | 1.64E-02 |
| TRANSLATION | Genes annotated by the GO term GO:0006412. The chemical reactions and pathways resulting in the formation of a protein. This is a ribosome-mediated process in which the information in messenger RNA (mRNA) is used to specify the sequence of amino acids in the protein. | 5 | 180 | 0.0278 | 4.63E-05 | 1.64E-02 |
| REGULATION_OF_TRANSLATION | Genes annotated by the GO term GO:0006417. Any process that modulates the frequency, rate or extent of the chemical reactions and pathways resulting in the formation of proteins by the translation of mRNA. | 4 | 93 | 0.043 | 5.11E-05 | 1.64E-02 |
| CELLULAR_BIOSYNTHETIC_PROCESS | Genes annotated by the GO term GO:0044249. The chemical reactions and pathways resulting in the formation of substances, carried out by individual cells. | 6 | 321 | 0.0187 | 7.20E-05 | 1.98E-02 |
| MULTICELLULAR_ORGANISMAL_DEVELOPMENT | Genes annotated by the GO term GO:0007275. The biological process whose specific outcome is the progression of an organism over time from an initial condition (e.g. a zygote or a young adult) to a later condition (e.g. a multicellular animal or an aged adult). | 10 | 1049 | 0.0095 | 9.16E-05 | 2.08E-02 |
| SYSTEM_DEVELOPMENT | Genes annotated by the GO term GO:0048731. The process whose specific outcome is the progression of an organismal system over time, from its formation to the mature structure. A system is a regularly interacting or interdependent group of organs or tissues that work together to carry out a given biological process. | 9 | 861 | 0.0105 | 1.05E-04 | 2.08E-02 |
| REGULATION_OF_GENE_EXPRESSION | the frequency, rate or extent of gene expression. Gene expression is the process in which a gene's coding sequence is converted into a mature gene product or products (proteins or RNA). This includes the production of an RNA transcript as well as any processing to produce a mature RNA product or an mRNA (for protein-coding genes) and the translation of that mRNA into protein. Some protein processing events may be included when they are required to form an active form of a product from an inactive precursor form. | 8 | 673 | 0.0119 | 1.08E-04 | 2.08E-02 |

**d** RNA-seq Symmetric assumption

| Gene Set Name | Description | # Genes in Overlap (k) | # Genes in Gene Set (K) | k/K | p-value | FDR q-value |
|---|---|---|---|---|---|---|
| REACTOME_CELL_JUNCTION_ORGANIZATION | Genes involved in Cell junction organization | 6 | 78 | 0.0769 | 1.83E-08 | 3.51E-05 |
| REACTOME_CELL_CELL_COMMUNICATION | Genes involved in Cell-Cell communication | 6 | 120 | 0.05 | 2.42E-07 | 2.33E-04 |
| REACTOME_NEF_MEDIATES_DOWN_MODULATION_OF_CELL_SURFACE_RECEPTORS_BY_RECRUITING_THEM_TO_CLATHRIN_ADAPTERS | Genes involved in Nef-mediates down modulation of cell surface receptors by recruiting them to clathrin adapters | 3 | 21 | 0.1429 | 1.22E-05 | 7.80E-03 |
| REACTOME_ADAPTIVE_IMMUNE_SYSTEM | Genes involved in Adaptive Immune System | 8 | 539 | 0.0148 | 2.12E-05 | 1.02E-02 |
| REACTOME_THE_ROLE_OF_NEF_IN_HIV1_REPLICATION_AND_DISEASE_PATHOGENESIS | Genes involved in The role of Nef in HIV-1 replication and disease pathogenesis | 3 | 28 | 0.1071 | 2.96E-05 | 1.14E-02 |
| REACTOME_MHC_CLASS_II_ANTIGEN_PRESENTATION | Genes involved in MHC class II antigen presentation | 4 | 91 | 0.044 | 4.51E-05 | 1.19E-02 |
| KEGG_ENDOCYTOSIS | Endocytosis | 5 | 183 | 0.0273 | 4.77E-05 | 1.19E-02 |
| REACTOME_SIGNAL_TRANSDUCTION_BY_L1 | Genes involved in Signal transduction by L1 | 3 | 34 | 0.0882 | 5.36E-05 | 1.19E-02 |
| HALLMARK_PROTEIN_SECRETION | Genes involved in protein secretion pathway. | 4 | 96 | 0.0417 | 5.56E-05 | 1.19E-02 |
| HALLMARK_APICAL_JUNCTION | Genes encoding components of apical junction complex. | 5 | 200 | 0.025 | 7.26E-05 | 1.40E-02 |

**Table S7. Functional enrichment analyses for genes whose expression distributions are associated with significant patient survival time for the TCGA OVC cohort.** Using the Investigate Gene Sets tool from GSEA/MSigDB, we detected significantly enriched terms and pathways for the top 100 most significant in survival time genes using the shape assumption (**a.** Microarray and **c.** RNA-seq), and top 100 most significant in survival time genes using a regular symmetric assumption (**b.** Microarray and **d.** RNA-seq). We used a threshold of FDR q-value < 0.05 or top 10 (whichever is less) was applied, and we computed overlaps with the following sets, H: Hallmark gene sets, C2 : KEGG and REACTOME, C5 : GO biological process, C6 : oncogenic signatures. Blue cells denote terms that were observed for both shape and symmetric assumptions, yellow cells denote terms that were unique to the shape or symmetric assumption (comparison between **a.** and **b.** and between **c.** and **d.**).

**a** Microarray Shape assumption

| Gene Set Name | Description | # Genes in Overlap (k) | # Genes in Gene Set (K) | k/K | p-value | FDR q-value |
|---|---|---|---|---|---|---|
| BIOPOLYMER_METABOLIC_PROCESS | Genes annotated by the GO term GO:0043283. The chemical reactions and pathways involving biopolymers, long, repeating chains of monomers found in nature e.g. polysaccharides and proteins. | 20 | 1684 | 0.0119 | 2.11E-10 | 4.07E-07 |
| POST_TRANSLATIONAL_PROTEIN_MODIFICATION | Genes annotated by the GO term GO:0043687. The covalent alteration of one or more amino acids occurring in a protein after the protein has been completely translated and released from the ribosome. | 11 | 476 | 0.0231 | 4.69E-09 | 3.10E-06 |
| PROTEIN_METABOLIC_PROCESS | Genes annotated by the GO term GO:0019538. The chemical reactions and pathways involving a specific protein, rather than of proteins in general. Includes protein modification. | 16 | 1231 | 0.013 | 4.84E-09 | 3.10E-06 |
| CELLULAR_PROTEIN_METABOLIC_PROCESS | Genes annotated by the GO term GO:0044267. The chemical reactions and pathways involving a specific protein, rather than of proteins in general, occurring at the level of an individual cell. Includes protein modification. | 15 | 1117 | 0.0134 | 1.00E-08 | 4.54E-06 |
| CELLULAR_MACROMOLECULE_METABOLIC_PROCESS | Genes annotated by the GO term GO:0044260. The chemical reactions and pathways involving macromolecules, large molecules including proteins, nucleic acids and carbohydrates, as carried out by individual cells. | 15 | 1131 | 0.0133 | 1.18E-08 | 4.54E-06 |
| PROTEIN_MODIFICATION_PROCESS | Genes annotated by the GO term GO:0006464. The covalent alteration of one or more amino acids occurring in proteins, peptides and nascent polypeptides (co-translational, post-translational modifications). Includes the modification of charged tRNAs that are destined to occur in a protein (pre-translation modification). | 11 | 631 | 0.0174 | 8.26E-08 | 2.65E-05 |
| BIOPOLYMER_MODIFICATION | Genes annotated by the GO term GO:0043412. The covalent alteration of one or more monomeric units in a polypeptide, polynucleotide, polysaccharide, or other biological polymer, resulting in a change in its properties. | 11 | 650 | 0.0169 | 1.11E-07 | 3.05E-05 |
| HALLMARK_INTERFERON_GAMMA_RESPONSE | Genes up-regulated in response to IFNG [GeneID=3458]. | 7 | 200 | 0.035 | 2.13E-07 | 5.12E-05 |
| SIGNAL_TRANSDUCTION | Genes annotated by the GO term GO:0007165. The cascade of processes by which a signal interacts with a receptor, causing a change in the level or activity of a second messenger or other downstream target, and ultimately effecting a change in the functioning of the cell. | 14 | 1634 | 0.0086 | 6.76E-06 | 1.44E-03 |
| PHOSPHORYLATION | Genes annotated by the GO term GO:0016310. The process of introducing a phosphate group into a molecule, usually with the formation of a phosphoric ester, a phosphoric anhydride or a phosphoric amide. | 6 | 313 | 0.0192 | 4.97E-05 | 8.96E-03 |

**b** Microarray Symmetric assumption

| Gene Set Name | Description | # Genes in Overlap (k) | # Genes in Gene Set (K) | k/K | p-value | FDR q-value |
|---|---|---|---|---|---|---|
| NUCLEOBASENUCLEOSIDENUCLEOTIDE_AND_NUCLEIC_ACID_METABOLIC_PROCESS | Genes annotated by the GO term GO:0006139. The chemical reactions and pathways involving nucleobases, nucleosides, nucleotides and nucleic acids. | 15 | 1244 | 0.0121 | 3.07E-08 | 4.24E-05 |
| BIOPOLYMER_METABOLIC_PROCESS | Genes annotated by the GO term GO:0043283. The chemical reactions and pathways involving biopolymers, long, repeating chains of monomers found in nature e.g. polysaccharides and proteins. | 17 | 1684 | 0.0101 | 4.41E-08 | 4.24E-05 |
| IMMUNE_SYSTEM_PROCESS | Genes annotated by the GO term GO:0002376. Any process involved in the development or functioning of the immune system, an organismal system for calibrated responses to potential internal or invasive threats. | 8 | 332 | 0.0241 | 4.08E-07 | 2.61E-04 |
| DNA_METABOLIC_PROCESS | Genes annotated by the GO term GO:0006259. The chemical reactions and pathways involving DNA, deoxyribonucleic acid, one of the two main types of nucleic acid, consisting of a long, unbranched macromolecule formed from one, or more commonly, two, strands of linked deoxyribonucleotides. | 7 | 257 | 0.0272 | 9.91E-07 | 4.77E-04 |
| ATF2_S_UP.V1_DN | Genes down-regulated in myometrial cells over-expressing a shortened splice form of ATF2 [Gene ID=1386] gene. | 6 | 187 | 0.0321 | 2.38E-06 | 9.16E-04 |
| HALLMARK_ALLOGRAFT_REJECTION | Genes up-regulated during transplant rejection. | 6 | 200 | 0.03 | 3.51E-06 | 1.12E-03 |
| ANATOMICAL_STRUCTURE_DEVELOPMENT | Genes annotated by the GO term GO:0048856. The biological process whose specific outcome is the progression of an anatomical structure from an initial condition to its mature state. This process begins with the formation of the structure and ends with the mature structure, whatever form that may be including its natural destruction. An anatomical structure is any biological entity that occupies space and is distinguished from its surroundings. Anatomical structures can be macroscopic such as a carpel, or microscopic such as an acrosome. | 11 | 1013 | 0.0109 | 6.65E-06 | 1.83E-03 |
| SYSTEM_DEVELOPMENT | Genes annotated by the GO term GO:0048731. The process whose specific outcome is the progression of an organismal system over time, from its formation to the mature structure. A system is a regularly interacting or interdependent group of organs or tissues that work together to carry out a given biological process. | 10 | 861 | 0.0116 | 1.01E-05 | 2.42E-03 |
| HEMOPOIETIC_OR_LYMPHOID_ORGAN_DEVELOPMENT | Genes annotated by the GO term GO:0048534. The process whose specific outcome is the progression of any organ involved in hemopoiesis or lymphoid cell activation over time, from its formation to the mature structure. Such development includes differentiation of resident cell types (stromal cells) and of migratory cell types dependent on the unique microenvironment afforded by the organ for their proper differentiation. | 4 | 77 | 0.0519 | 1.90E-05 | 4.06E-03 |
| IMMUNE_SYSTEM_DEVELOPMENT | Genes annotated by the GO term GO:0002520. The process whose specific outcome is the progression of an organismal system whose objective is to provide calibrated responses by an organism to a potential internal or invasive threat, over time, from its formation to the mature structure. A system is a regularly interacting or interdependent group of organs or tissues that work together to carry out a given biological process. | 4 | 81 | 0.0494 | 2.32E-05 | 4.46E-03 |

**c** RNA-seq Shape assumption

| Gene Set Name | Description | # Genes in Overlap (k) | # Genes in Gene Set (K) | k/K | p-value | FDR q-value |
|---|---|---|---|---|---|---|
| REACTOME_MICRORNA_MIRNA_BIOGENESIS | Genes involved in MicroRNA (miRNA) Biogenesis | 3 | 23 | 0.1304 | 1.66E-05 | 2.34E-02 |
| REACTOME_REGULATORY_RNA_PATHWAYS | Genes involved in Regulatory RNA pathways | 3 | 26 | 0.1154 | 2.43E-05 | 2.34E-02 |
| HALLMARK_EPITHELIAL_MESENCHYMAL_TRANSITION | Genes defining epithelial-mesenchymal transition, as in wound healing, fibrosis and metastasis. | 5 | 200 | 0.025 | 7.62E-05 | 3.67E-02 |
| HALLMARK_MTORC1_SIGNALING | Genes up-regulated through activation of mTORC1 complex. | 5 | 200 | 0.025 | 7.62E-05 | 3.67E-02 |

**d** RNA-seq Symmetric assumption

| Gene Set Name | Description | # Genes in Overlap (k) | # Genes in Gene Set (K) | k/K | p-value | FDR q-value |
|---|---|---|---|---|---|---|
| NEGATIVE_REGULATION_OF_TRANSFERASE_ACTIVITY | Genes annotated by the GO term GO:0051348. Any process that stops or reduces the rate of transferase activity, the catalysis of the transfer of a group, e.g. a methyl group, glycosyl group, acyl group, phosphorus-containing, or other groups, from a donor compound to an acceptor. | 4 | 35 | 0.1143 | 1.01E-06 | 1.94E-03 |
| INACTIVATION_OF_MAPK_ACTIVITY | Genes annotated by the GO term GO:0000188. Any process that terminates the activity of the active enzyme MAP kinase. | 3 | 14 | 0.2143 | 3.47E-06 | 3.34E-03 |
| NEGATIVE_REGULATION_OF_MAP_KINASE_ACTIVITY | Genes annotated by the GO term GO:0043407. Any process that stops, prevents or reduces the frequency, rate or extent of MAP kinase activity. | 3 | 17 | 0.1765 | 6.45E-06 | 4.14E-03 |
| SIGNAL_TRANSDUCTION | Genes annotated by the GO term GO:0007165. The cascade of processes by which a signal interacts with a receptor, causing a change in the level or activity of a second messenger or other downstream target, and ultimately effecting a change in the functioning of the cell. | 14 | 1634 | 0.0086 | 1.10E-05 | 5.29E-03 |
| NEGATIVE_REGULATION_OF_CATALYTIC_ACTIVITY | Genes annotated by the GO term GO:0043086. Any process that stops or reduces the activity of an enzyme. | 4 | 69 | 0.058 | 1.57E-05 | 6.05E-03 |
| ATF2_S_UP.V1_DN | Genes down-regulated in myometrial cells over-expressing a shortened splice form of ATF2 [Gene ID=1386] gene. | 5 | 187 | 0.0267 | 5.55E-05 | 1.78E-02 |

**Table S8. Overlap between genes that were previously identified in a prognostic signature and those appearing in our list of genes with different expression distributions for the TCGA AML RNA-seq data set.** No overlap was found with the TCGA AML microarray dataset.

| | Prognostic Signatures |
|---|---|

| **Gene Expression Distributions** | Valk et al. (2004)[1] | Bartholdy et al. (2014)[2] | Marcucci et al. (2011)[3] | Gentles et al. (2010)[4] | Eppert et al. (2011)[5] Phenotypic Signature | Eppert et al. (2011)[5] Leukemia Stem Cell Signature (LSC-R) | Eppert et al. (2011)[5] Hematopoietic Stem Cell Signature (HSC-R) | Li et al. (2013)[6] |
|---|---|---|---|---|---|---|---|---|
| **Normal** | - | RNASE1 | - | LTB | - | - | - | - |
| **Lognormal** | - | - | - | - | - | - | - | - |
| **Bimodal** | - | - | - | - | - | - | - | - |
| **Cauchy** | - | - | - | - | - | - | - | - |
| **Gamma** | - | TLK2 | - | - | - | - | - | - |
| **Total Overlap** | 0 | 2 | 0 | 1 | 0 | 0 | 0 | 0 |
| **Number of Genes in Signature** | 175 | 561 | 17 | 50 | 47 | 47 | 143 | 24 |
